# Causal Understanding of the Stone Dropping Task in Two Species of Macaw

**DOI:** 10.1101/2020.08.24.264390

**Authors:** Laurie O’Neill, Anthony Picaud, Ronan Hastings, Nina Buffenoir, Manfred Gahr, Auguste M.P. von Bayern

## Abstract

Causal understanding in animal cognition can be divided into two broad categories (Woodward, 2011): learned associations between cause and effect (Le Pelley et al., 2017) and understanding based on underlying mechanisms (Johnson and Ahn, 2017). One experiment that gives insight to animals’ use of causal mechanisms is the stone-dropping task. In this, subjects are given an opportunity to push a platform to make it collapse and are then required to innovate dropping a stone tool to recreate the platform collapsing (von Bayern et al., 2009). We describe how 16/18 subjects of two species of macaw (*n=18; Ara ambiguus (n=9)* & *Ara glaucogularis (n=9)*) were able to innovate the solution in this task. Many of the subjects were able to innovate the behaviour through exploratory object combination, but it is also possible that a mechanistic understanding of the necessity for contact with the platform influenced some subjects’ behaviour. All the successful subjects were able to recreate their novel stone-dropping behaviour in the first or second trial after innovation (and all trials thereafter) and they were also able to do the behaviour increasingly faster. This suggests they also rely on learned associations of cause and effect. However, in a transfer task in which subjects had to guide a stick tool to make it touch a differently positioned platform, all but one of the subjects failed. This would suggest that the majority of the subjects were not using an understanding of platform contact to solve the task, although the subjects’ difficulty with using stick tools may have also affected their performance in this transfer.

## Introduction

Cognition is the process by which an animal mediates between incoming senses and outgoing behaviour. Causal cognition is specifically the individual’s understanding of the cause and effect relationships they observe in their environment. Some have suggested that causal reasoning skills are a key tenet of complex cognition (Emery and Clayton, 2004) and thus are likely to be present in multiple species that have evolutionary pressures for ‘intelligent’ behaviour (Van Horik et al., 2012). It could be that it is one of the cognitively complex processes that make some species more behaviourally flexible, innovative and thus successful at rapidly adapting to new environments (Lefebvre et al., 1997; Sol et al., 2007, 2005, 2002). By studying how different species mentally represent causal relationships we can learn more about the evolution of causal understanding and its role in the evolution of complex cognition.

Causal relationships are probably grasped through more than one cognitive process. As an example, think of a crow that opens a snail shell by dropping it onto a concrete path running through its territory. Does the crow only recognize the spatial connection between the successful opening of shells and the concrete area or do they specifically recognize that the concreted area is made of a harder material, and that hard materials break things more easily. Within this example is the description of two forms of causal understanding. The former form of understanding is an associatively learnt account of a causal link between events that are spatially or temporally connected (Hanus, 2016; Le Pelley et al., 2017) and has been referred to as ‘difference making’ understanding (Woodward, 2011). The latter form of understanding specifies the underlying causal mechanism, for example the physical forces that link a cause to its effect (Johnson and Ahn, 2017) and has been referred to as ‘geometrical-mechanical’ understanding (Woodward, 2011). Studies investigating a species’ causal cognition do not always explicitly define causal understanding between these two types. However, they are often recognized under different names. For example, geometric-causal bears similarities to ‘folk-physics’ (Povinelli, 2000) whereas difference-making understanding is regularly dismissed as not being causal understanding due to its associatively-learnt properties. Although difference-making understanding requires learning, a learned causal link is still a form of causal cognition (Hanus, 2016; Le Pelley et al., 2017)

Studies on different parrot species suggest they are capable of many types of complex and flexible physical problem solving skills (Lambert et al., 2018). They also one of the groups of animals that have both relatively large, neuronally dense brains (Gutiérrez-Ibáñez et al., 2018; Herculano-Houzel, 2017; Iwaniuk et al., 2005; Olkowicz et al., 2016) and known environmental stochasticity (Toft et al., 2016) that suggest they may use complex causal cognition processes in their daily life (Osvath et al., 2014; Van Horik et al., 2012). A notable example of their technical competence is their ability to use tools in captivity. In the wild, there are just a few examples of parrots using tools (Goodman et al., 2018; Heinsohn et al., 2017; Osuna-Mascaró and Auersperg, 2018; Villegas-Retana and Araya-H., 2017; Wood, 1984), but in captivity two species, kea (*Nestor notabalis*) and Goffin’s cockatoos (*Cacatua goffiniana*), have been shown to innovate stick tool use in problem-solving situations (Auersperg et al., 2011; Auersperg et al., 2012). The tool use innovation reported from Goffin’s cockatoos was particularly remarkable as one individual even innovated the manufacture of a tool. This kind of flexible tool-use from a non-habitually tool using animal suggests the species exhibits elements of complex physical understanding of object properties (Bird and Emery, 2009; Kacelnik, 2009). It is thus important to test the causal understanding and cognitive abilities of more parrot species on physical causal understanding tasks to learn about its role in all of parrots’ cognitive evolution.

The species tested here belong to the genus *Ara* (the macaws) in the parrot superfamily *Psittacoidea*. They are relatively distantly related to the Kea (superfamily *Strigopoidea*) and the Goffin’s cockatoos (superfamily Cacatuoidea) with a last common ancestor approximately 50mya (Kumar et al., 2017; Wright et al., 2008). We tested two species of macaws: *Ara glaucogularis* and *Ara ambiguus*. These macaws had previously completed a range of causal understanding tasks as part of a larger cognitive test battery (Krasheninnikova et al., 2019) and also completed a modified trap-tube task (O’Neill et al., 2018). Both of these experiments were object choice tasks with two or three options to choose from and it turned out the subjects developed some rules of thumb (heuristics), such as side biases or random choice, that allowed subjects to attain a satisfactory amount of rewards without any ‘cognitive effort’. To avoid this, we decided to design an experiment that did not include choice elements but instead presented the subjects with a novel problem-solving task.

We gave the subjects the stone-dropping task (Bird and Emery, 2009; von Bayern et al., 2009). In this task, a subject was first able to push a collapsible platform with their beak to release a food reward. The platform was then placed out of reach to see if the subject could recreate the effect of the platform pushing behaviour by innovating a novel causal input: dropping a stone onto it. Two out of four New Caledonian Crows (*Corvus moneduloides*) were able to recreate this force by dropping a stone tool onto the platform (von Bayern et al., 2009). It is not clear exactly what kind of understanding these crows used to solve the task. They may have understood that they were required to make contact with the platform and had recognised that dropping a stone was an alternative form of contact. The understanding that contact was the key factor would suggest that the crows had an understanding based on the mechanism underlying the function of the apparatus. However, it appears unlikely that they understood more complex mechanical properties of the task such as the importance of the stone’s weight, as other New Caledonian crows did not initially discriminate between dropping heavy and light stones in a follow up task from a replication study (Neilands et al., 2016). However in this replication, only a single New Caledonian crow of twelve was able to innovate the stone-dropping behaviour. Further, it was interesting to note that these subjects were also able to learn that only heavy stones were functional in making the platform collapse (Neilands et al., 2016).

It seems unlikely that the birds’ initial stone-dropping actions were down to a learned, difference-making, causal understanding, as they had never observed the effects of a stone being dropped onto the platform before. However, many subjects began to drop more stones into the apparatus after the platform had collapsed, suggesting that they had also formed a difference-making understanding that the dropping of stones led to the appearance of the reward (Neilands et al., 2016). It was unlikely that the behaviour was driven by spontaneous exploration as the birds were given extensive opportunity to explore the apparatus in the presence of stones, before they were given the opportunity to push the platform with their beak, and they never dropped a stone in at this point (von Bayern et al., 2009).

The rationale of the current experiment was thus as follows. Subjects are given an opportunity to solve the task without any experience of how the platform in the apparatus can move. This phase is called the pre-test. If subjects are able to solve the experiment at this stage (see supplementary video 1 for an example of a successful solution) then it suggests they are capable of solving the task through exploratory behaviour alone. This is because they cannot be using a causal understanding of how an apparatus works if they have never had experience of any of its functional properties before.

They were then given an opportunity to experience the collapsibility of the platform and resulting release of food by being allowed to collapse it themselves directly by pushing it (see supplementary video 2). This platform pushing experience gives the subjects knowledge of the functional mechanism underlying the required action of the task, but not the direct actions they need to take to solve the task by dropping stones. Following this, another test phase is given (critical test phase), which is identical to the pre-test phase except it was now possible to see if and how experience of the functional mechanisms had changed their behaviour. If they immediately began to drop rocks on the platform at this stage, it suggested the subjects had developed an understanding of the causal mechanism of the platform, i.e. that it can be collapsed by exerting force or making contact with it, and were trying to activate this mechanism in another way.

After this, it is vital to then see if subjects continue to persist with the stone-dropping behaviour. This is to ensure that they have not done it accidentally. It is also possible to then see the extent to which the subjects recognised the effects of their own behaviour from the first stone drop onwards. If they begin to drop stones with a reduced latency after the first successful stone drop, it suggests an egocentric understanding of the effects they have caused on the apparatus. Thus they can learn quickly to repeat the successful behaviour.

If the subjects were successful in dropping stones onto the platform, we also planned a transfer task to specifically observe whether they recognised the importance of contact between an object and the platform to make it collapse. This transfer task involved partially moving the platform from directly underneath the tube into which they could drop stones in, this meant objects dropped into the tube would not automatically make contact with the platform. Instead subjects would have to use a stick tool inserted at an angle to make contact with the platform. Thus subjects that only had a learned causal understanding that objects dropped into the apparatus made rewards appear would struggle to complete this transfer task. Success in this transfer task would add more confirmatory evidence that subjects recognised the mechanism that made the apparatus function.

We expected that at least some individuals of the two macaw species tested would be able to drop stones. They have shown previous success in another test for causal understanding (O’Neill et al., 2018) and they are known to be highly exploratory and keen at handling objects, so were likely to interact with the stones given to them. We were unsure if the macaws would do this immediately after the experience phase given to them, as they have not previously shown mechanistic causal understanding in previous experiments.

## Methods

### Subjects and housing

Nine *A.ambiguus* and nine *A.glaucogularis* were tested. Their age and sex are shown in Table 1. All the birds were hand-raised and group reared by the Loro Parque Foundation in Tenerife, Spain, and all were housed in the Comparative Cognition Research Station, within the Loro Parque zoo in Puerto de la Cruz, Tenerife. The birds were housed in groups of 2-8 individuals, according to species and age, in seven aviaries. Six of these aviaries were 1.8 x 6 x 3 metres (width x length x height) and one was 2 x 6 x 3 metres. Windows of 1 x 1 m could be opened between the aviaries to connect them together. One half of the aviaries was outside, so the birds followed the natural heat and light schedule of Tenerife. The half inside the research station and was lit with Arcadia Zoo Bars (Arcadia 54W freshwater Pro and Arcadia 54W D3 Reptile Lamp) that followed the natural light schedule.

**Table 1.**
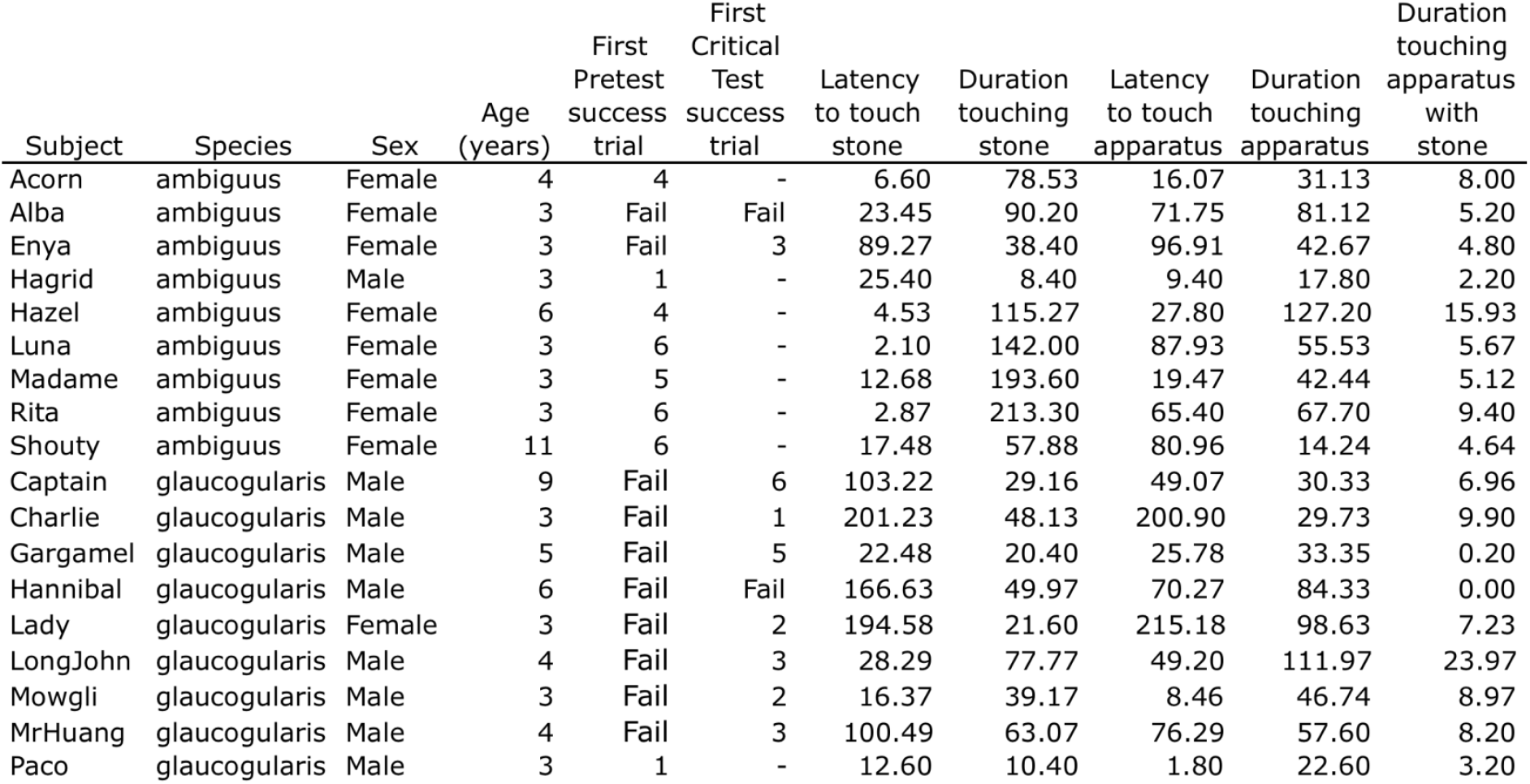
Description of subjects and their success in the stone dropping task. ‘Fail’ indicates that the subjects completed six sessions of the given phase without reaching the successful criteria. Numbers indicate the session in which subjects had their first successful trial. Dashes indicate that the subject did not do the phase because they had already succeeded in the experiment. The five columns on the right indicate individuals average exploration times (in seconds) in trials up until, and including, their first successful trial.

A large variety of fresh fruit and vegetables were provided first thing in the morning and again in the afternoon. In the afternoon, each individual received a portion of Versele Laga Ara seed-mix that was modified based on their daily weight. The parrots were never starved, controlled portions of seeds were fed to prevent overeating and obesity. Testing took place at least an hour after the parrots had breakfast to ensure there was sufficient motivation to get food rewards. Unless otherwise specified, walnut halves were used as rewards. These were highly prized by all subjects, and they were able to obtain them on a daily basis through voluntary testing.

### Experimental rooms

Experiments took place in separate testing rooms away from the aviaries. All birds had been previously trained to enter in them. These rooms were 1.5 x 1.5 x 1.5 m and also lit with Arcadia Zoo Bars to cover the birds’ full visual range. One wall of each one of the experimental rooms had a 50 x 25 cm window through which an experimenter could place apparatuses into the testing room from a neighbouring chamber. This window could be occluded with a white board so that the experimenter could hide anything they were doing from the subjects’ view, such as re-setting apparatuses between trials. The experimenter always wore mirrored sunglasses during experiments to prevent their gaze being a cue for the parrots. A second wall was made of sound proofed one-way glass so that zoo visitors could observe experiments without disturbing the subjects.

### Apparatus

The apparatus had a hinged platform held up by two 8×8mm neodymium disc magnets (www.supermagnete.de, 2.5kg strength) that magnetised to two screws. A transparent acrylic shell (25 x 20 x 10.5 cm), marked with blue stripes to show the solidity of the acrylic, enclosed the platform. A reward was placed on top of this platform and was inaccessible unless the magnets released the hinged platform into a sloped position, allowing the reward to roll out from another hole at the bottom of the apparatus (Figure 1). For the pre-test and critical test, the only access hole to the platform was via a small rectangular tube, placed on top of the shell. The platform was 10 cm below the top of this tube, which meant the parrots could not reach in and touch the platform. The opening at the top of the tube was a rectangle of 7 x 4.5 cm, which was also small enough to prevent the birds from placing their heads directly inside the hole. The entire apparatus was mounted on a solid wooden board (45 x 30 x 2.5 cm).

**Figure 1.**
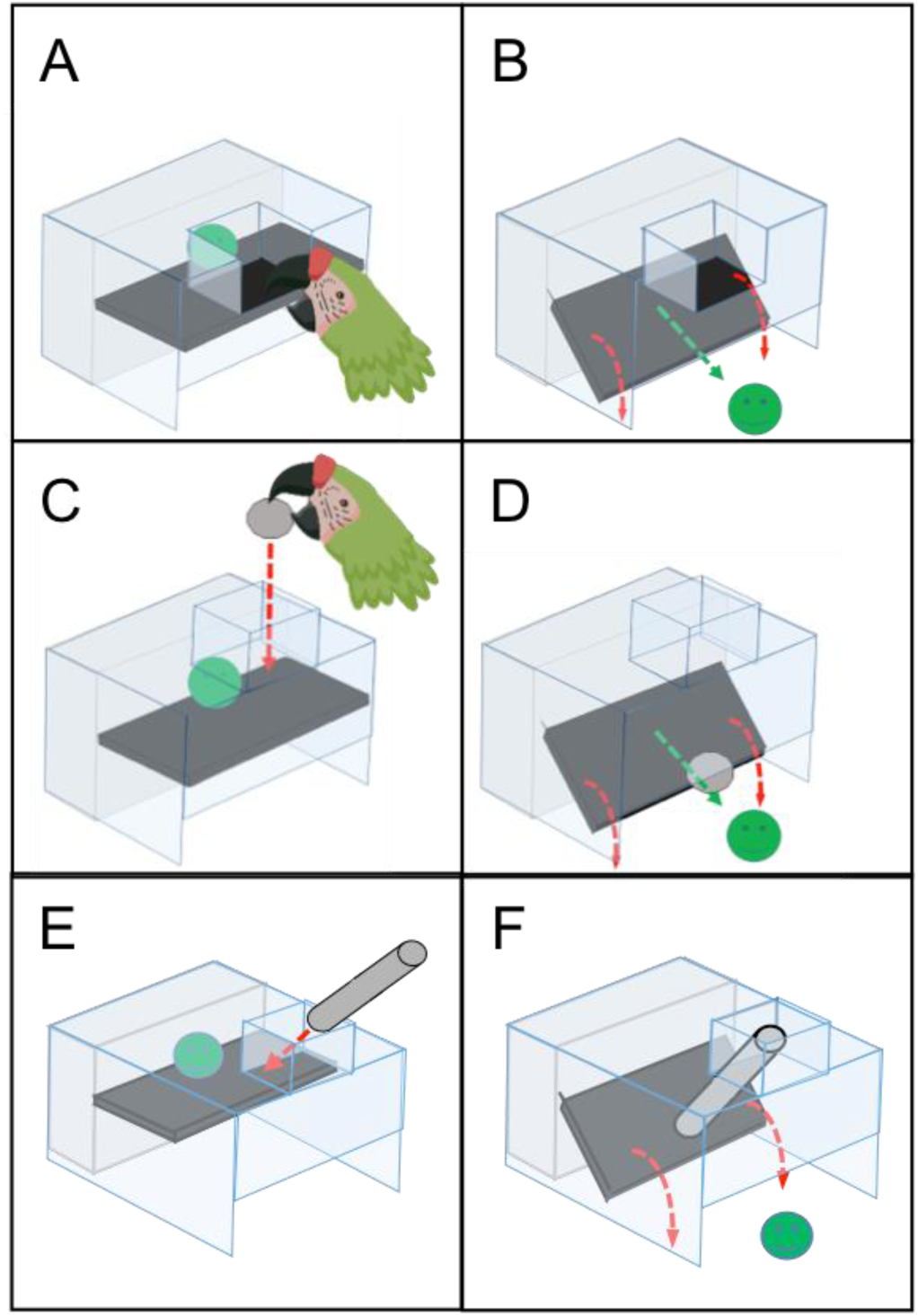
All of the apparatus used in the stone dropping test, the platform pushing experience and the stick dropping transfer. A&B show the apparatus setup for the platform pushing experience. In this stage, subjects could push the black platform (A) causing it to detach from a pair of magnets and allowing a reward to fall out the bottom (B). C&D show the apparatus setup for the pre-test and critical test phases of stone dropping task, which occurred before and after the platform pushing experience respectively. In these stages, the subjects could only make the platform collapse by dropping a stone through the tube in the top of the apparatus (c), to release a reward (D). E&F show the stick transfer task, in which the transparent acrylic shell was extended so that the tube was no longer directly above the platform. This meant that a stick had to be inserted at an angle into the tube (E) so that it touched the platform (F).

A slightly modified version of the apparatus was used for the platform-pushing phase. A different transparent acrylic shell was placed on the apparatus that allowed the subjects to access the platform. In place of the tube, this shell had an opening so that the birds could push the platform directly with their beaks (Figure 1, A & B). However, this structure still blocked the parrots’ direct access to the reward that was resting on top of the platform.

The stone tools provided in the pre-test and critical tests were natural volcanic rocks taken from the beach in Puerto de la Cruz, Tenerife. They were returned to the beach after testing was finished. They had a diameter of 4.5-5.5 cm and weighed between 60-100 g. It was ensured they were all sufficiently heavy to break the connection between the magnets holding up the platform when dropped from a height of 5 cm, i.e. half the height of the tube.

### Experimental procedures

Within these descriptions, a ‘session’ is a single period of time in which a subject was taken to a testing room, a ‘trial’ is a single opportunity to interact with the test apparatus and obtain a reward. Thus multiple trials could take place within a single session. A single trial could last for a maximum of ten minutes; if a subject had not succeeded within ten minutes then the trial *and* the session ended. If a subject succeeded within ten minutes, then the apparatus was rebaited and subjects were given another trial. There was a maximum of six trials in a session. Hypothetically sessions had a maximum time of thirty minutes, so as not to over-test the subjects, but this time limit was never reached. Typically subjects only had one session of testing per day. All subjects individually took less than a month to complete the whole experiment, from habituation to final critical test. The one exception was Hannibal, who took two months. Testing of the birds began in May 2017 and finished in January 2018.

### Habituation

Habituation to the apparatus and the stone tools was necessary to ensure the subjects were not scared of either of these and thus, to exclude neophobia as a confounding factor. Although parrots are typically quite neophilic (Auersperg et al., 2015; Mettke-Hofmann et al., 2002), so are unlikely to have a neophobic response to novel apparatus and objects, there is variation between individuals. Specifically, older parrots may be more likely to show neophobic responses (O’Hara et al., 2017). As the subjects were of different ages (see Table 1), it was important to ensure that all subjects were equally comfortable with the apparatus.

Every subject was first habituated to the apparatus without the necessary tools. These sessions took place in the experimental rooms, with an experimenter sitting in the neighbouring chamber. Once the subject was in the experimental room, the experimenter introduced the apparatus through the window. Subjects had five minutes to approach the apparatus and take a piece of walnut placed at the base of the apparatus (where the reward would fall out in a successful trial). If they took it, the experimenter waited for 30 seconds then removed the apparatus. Next, the experimenter, out of sight of the subject, rebaited the apparatus and reintroduced it a minute after removing it. The subjects had up to six of these trials per session and if they took six walnut pieces in a row within a single session, they moved onto testing.

In parallel, the subjects were habituated to the stone tools in their home aviaries. The stones were placed inside the aviaries for the parrots to approach and explore as a group. Each group of birds approached, picked up, and explored the tools extensively (although this was not specifically measured). Additionally, each individual was also given an opportunity to explore the tools whilst alone in an isolated area of the aviary. Each individual had held and interacted with the stones more than once before they started testing.

### Pre-test and Critical test procedure

The pre-test and critical test procedure were identical in both setup and procedure as detailed below. However, the birds entered the critical test phase only after failing to find a solution in the pre-test and after they had undergone the platform pushing experience phase (see below), during which they had an opportunity to learn about the underlying mechanism of the apparatus.

The pre-test was implemented in order to examine whether the subjects would solve the task without any prior knowledge of the underlying platform mechanism. If subjects naively placed stones into the apparatus this would indicate that they did not need to employ causal reasoning processes, but that their solution could by explained by more simple means, such as a tendency to insert stones into cavities or to combine them with other objects regardless of the causal outcome of this behaviour. On the contrary, if they did not drop stones in the pre-test, but only began to do so in the critical test, this could be seen as an indication that having experience of the underlying causal mechanism of the task (i.e. the collapsibility of the platform; see platform pushing experience below) caused a change in the subjects’ behaviour, suggesting the subjects were capable of a more complex form of causal understanding based on mechanisms.

The protocol for the pre-test and critical test was as follows. Subjects were given six testing sessions once per day, with both the baited test apparatus and tools present. Each session had between one to six trials, depending on the subject’s success, i.e. they were given more trials if they succeeded in a trial until the six trial limit for a session was reached. Initially, the whole apparatus was prepared out of sight in a neighbouring chamber to the subject’s testing room, including baiting the apparatus with an attractive reward (half a walnut) and placing two stone tools on each side of the apparatus (four in total). The apparatus was then pushed into the subject’s room through the window. The subject was allowed to interact with the apparatus for ten minutes. If they were not successful in this time, the trial (and the session) ended, the apparatus was removed and the subject was returned to its social group. If, however, a subject was successful in this time-span, and they were able to make the platform collapse through dropping stones into the tube, they would be given further trials immediately within the same session in order to examine whether they would continue to succeed in subsequent trials. The repetition of trials immediately after a successful stone drop was vital for verifying if the subjects could replicate their success and thus had grasped the causal outcome from their specific action, i.e. dropping the stone into the tube, or whether they had succeeded accidentally without recognising the difference-making properties of their behaviour. If they immediately repeated the behaviour after doing it once, it suggested that the behaviour was either originally purposeful, or that it was initially accidental but the birds at least had capable egocentric understanding of their behaviour. To implement a replication trial, the experimenter waited 30 seconds after the subject had made the platform collapse and then removed the apparatus. They would re-bait it out of sight of the subject and replace it one minute after removal. From the moment the apparatus was placed back in the subject’s compartment, they had another ten minutes to solve the task again. If they were not successful in a repeat trial, the session was ended. The apparatus was re-baited up to five more times within a session (thus a maximum of six trials per session). Testing stopped either if a subject reached criterion, which meant that the subject obtained rewards 12 times in a row in two continuous sessions (6 successful trials per session), or they had six full, valid sessions with no successful trials. If subjects had their first successful trial in the sixth session of testing then they were given a seventh session to see if they would reach the successful criterion at this point.

We considered a trial as unsuccessful, but valid, if the subjects touched either the apparatus or stone tools at any point. Thus trials in which subjects did not approach the apparatus or tools were counted as invalid. Testing sessions also ended after these invalid trials.

### Platform pushing experience

If subjects were unsuccessful in the pre-test, they were given the platform pushing experience before they moved onto the critical test (supplementary video 2). Its purpose was to give the subjects the opportunity to learn about the functional mechanism of the platform, i.e. the collapsibility of the platform if force was exerted onto it, without providing any cues to the required problem solving behaviour if the target platform was out of reach, for example dropping a stone tool onto it. They had to push down the platform directly with their beak to release a reward. To this end, the apparatus had to be slightly modified to allow for this (Figure 1, A & B). The subjects learned to push down the platform in the following way.

To begin with subjects were first provided with one 10-minute session to establish if they would spontaneously push the platform. If they did, then it would be re-baited five more times in this first session. They would continue to have 10-minute experience sessions until they reach a criterion of pushing the platform twelve times in a row (with six trials a session).

If subjects did not push the platform spontaneously, they were gradually trained to push the platform via shaping. Firstly, the magnets holding up the platform were weakened by placing layers of tape between the magnet and the screws they magnetised to. Just touching the platform was now enough to make it collapse. We gave them motivation to touch the platform by placing a small reward (a single sunflower seed) on it. When they picked up this reward, the platform collapsed and released the larger walnut reward from inside the apparatus. If they succeeded, the apparatus was removed, re-baited and replaced five more times for the subjects to repeat this behaviour (six in total). In the next session, the full magnet strength was restored but the small reward was still placed in the pushing area. Six more successes were required to pass this stage. Then, in the final sessions, the subjects had to push the full strength platform without the small reward incentive. They had to do this 12 times in a row in two sessions (six trials per session) to finally complete the platform pushing experience and move on to the critical test phase (as described above).

### Stick Transfer

Subjects would only proceed to this task if they had successfully completed the stone-dropping task, in either the pre-test or the critical test. It used the same apparatus as the stone-dropping task except that the acrylic shell that enclosed the platform was moved 5cm further forward (Figure 1 E&F; supplementary video 4). This change meant that objects dropped into the tube on top of the apparatus would not automatically hit the platform. Instead, they had to insert a stick at an angle to make the end of the stick touch the platform. Dropping the sticks vertically would mean the sticks missed the platform, thus it wouldn’t collapse and release a reward.

As these birds had never used a stick tool before, it was unclear what kind of stick would be easier for them to use, so we provided them with two different options. Both options were 18cm long and made of aluminium, but one option was solid with a 0.8 cm diameter and the other option was hollow with a 2 cm diameter. The hollow stick was open at each end so that the macaws could place the end of their beak inside the stick; we thought this may have been an easier way for them to grasp the stick tool. To further make the stick tools easier to grip for the birds, they were wrapped with a single layer of silver duct tape. Additionally, multiple layers of black electrical tape were wrapped around each end of the sticks to make it harder for them to remove the duct tape from the sticks. Subjects were habituated to these sticks using the same protocol as used for the stone tools.

This task only consisted of a critical test phase as all the subjects that took part had experience of the mechanism of the platform from the stone-dropping task. Thus, the aim of this task was to evaluate if the birds could use the stick tool to produce the same result as in the previous task. The procedure followed for this task was the same as the pre-test/critical test method as the stone-dropping task.

### Behavioural coding

All experiments were recorded on four static CCTV cameras. These covered all angles of the testing room. Two recordings from separate cameras were saved for each experimental session, but more recordings were saved if it was necessary for specific trials, e.g. if there was partial occlusion of a camera view from the subject standing in the way. The following exploratory behaviours/variables were scored from the videos using Solomon coder (András Péter, solomon.andraspeter.com): the amount of time (in seconds) subjects spent touching the apparatus, the amount of time they spent touching the stick or stone tools and the amount of time they spent touching the apparatus with the tools. These exploratory behaviours were scored from the moment the apparatus entered the subjects testing chamber until they either succeeded in a trial or until they reached the time limit for the trial (failure). We also counted the number of times subjects placed the stone or stick tools onto the apparatus, without this leading to a successful solution. The tool was defined as ‘on top’ of the apparatus if the subject released the stone from its beak and it remained stationary on top of the apparatus after release, but was not dropped inside the apparatus (i.e. onto the platform). Furthermore we scored the latency to solve the task for each individual and their latency to complete the successful behaviour in each of their successful trials. This latency was calculated from the moment the apparatus was placed inside the testing chamber until the platform had collapsed. Finally, we scored the number of times subjects dropped the stone and stick tools inside of the apparatus in each trial. This final behaviour was counted from the time the apparatus was placed inside the testing room until the moment the apparatus was removed from the testing room, which included the 30 seconds after subjects had successfully dropped a stick or stone tool onto the platform to make it collapse.

## Results

16 out of 18 subjects successfully and consistently dropped stones onto the platform (Table 1; supplementary video 5 shows each individuals first innovation of the behaviour). The other two birds, one of each species, never dropped any stones. Of the birds that succeeded, eight reached the criterion in the pre-test phase and the other eight reached it in the critical test phase (Figure 2). Of the birds that reached the criterion in the pre-test phase, seven were *A. ambiguus* and one was an *A. glaucogularis*. Thus, those that succeeded in the critical test phase were one *A. ambiguus* and seven *A. glaucogularis*.

**Figure 2.**
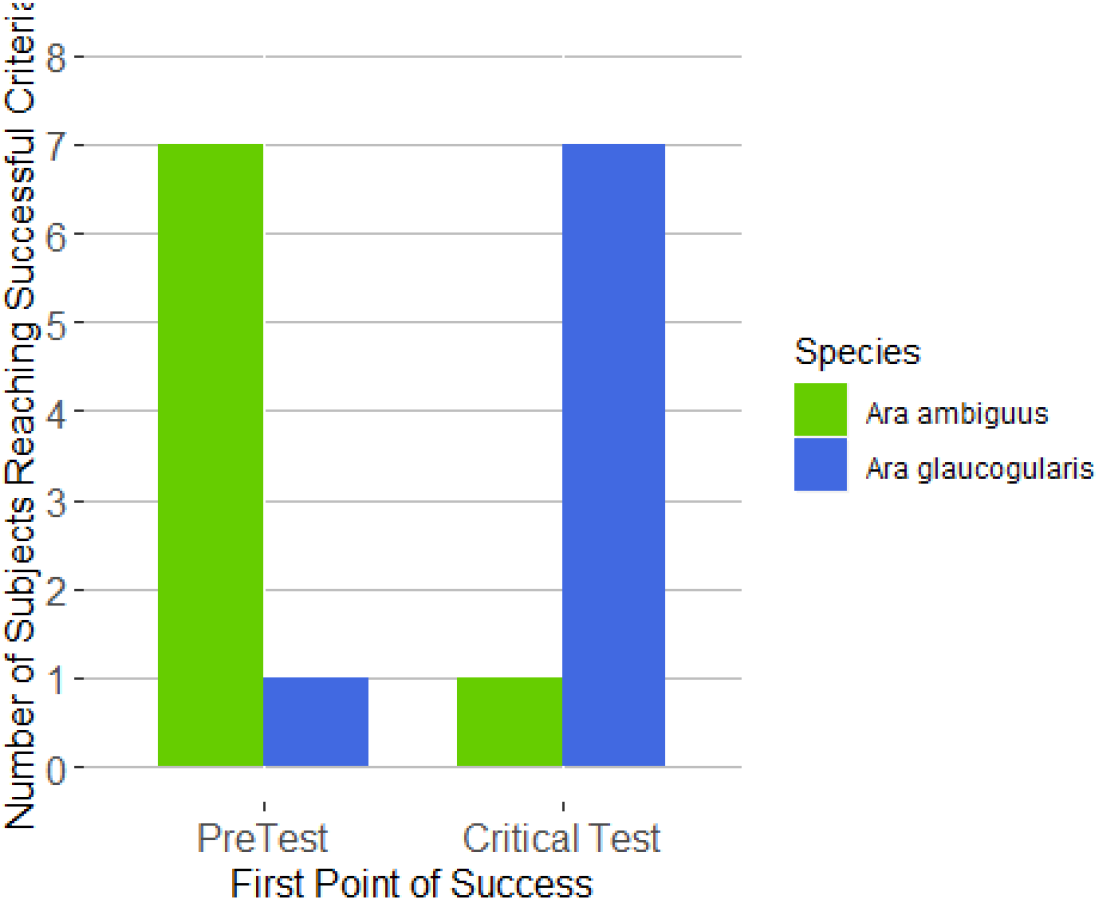
The Number of Subjects successfully passing the stone dropping task at each stage. Many birds were able to pass the stone dropping test from both species. However there were differences between the two species as to what stage they could pass the test. The majority of the Ara ambiguus were able to complete the task in the pre-test phase, whereas the majority of the Ara glaucogularis only completed the task in the Critical Test Phase. Two birds, one of each species, failed the task.

The *A. ambiguus* spent significantly more time interacting with the stone tools (mean = 104 seconds, sd = 69 seconds) than the *A. glaucogularis* (mean = 40 seconds, sd = 22 seconds) in all trials before their first successful stone drop (Figure 3; Welch’s two sample t-test, *t_9.6_* = 2.67, *p* = 0.02). There were no significant differences between the two species in the other exploratory measures; there was no difference between how much the two species interacted with the apparatus nor was there a difference in time they spent touching the apparatus with a tool held in their beak.

**Figure 3.**
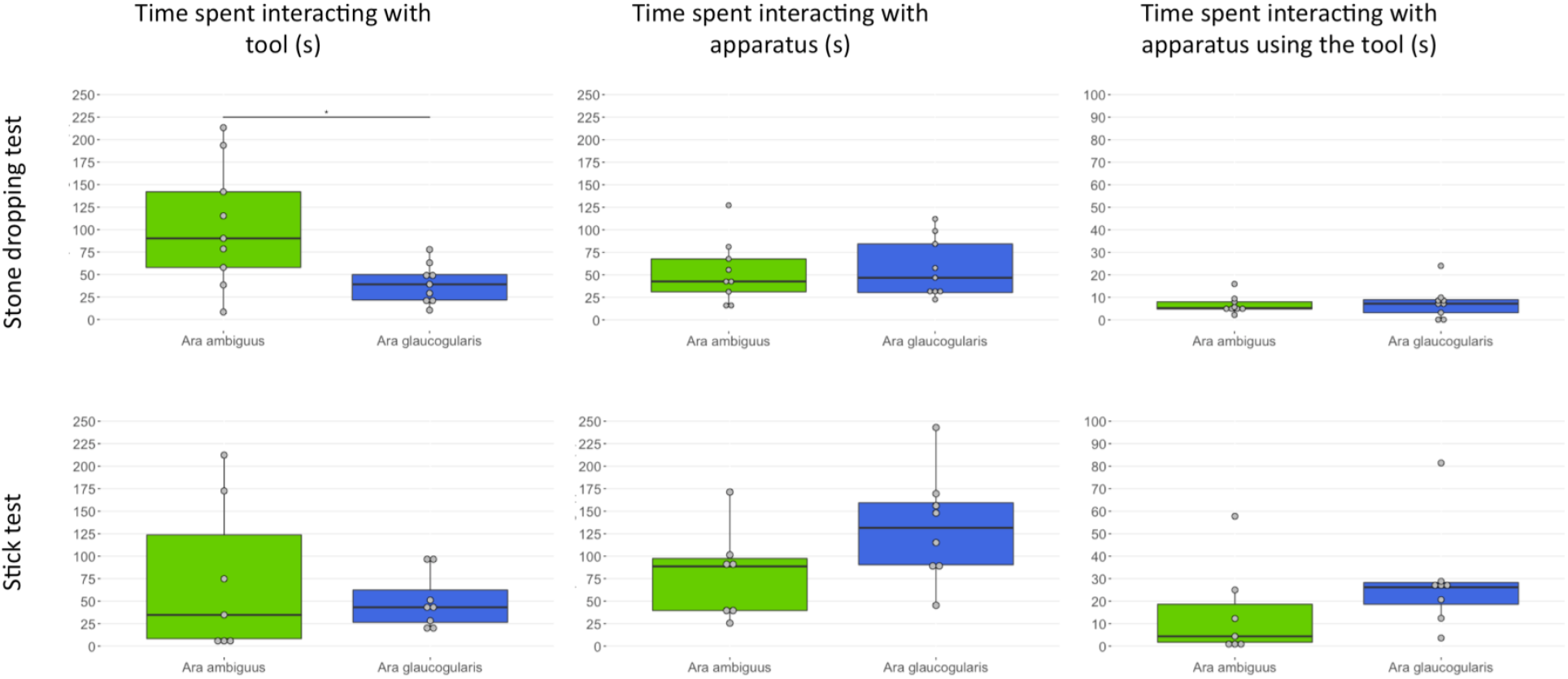
The average amount of time individuals spent interacting with the tools, the apparatus and a combination of both during both the initial stone dropping test and the stick test transfer. Only the trials prior to their first success are shown, as these are the only trials in which the birds were ‘exploring’, thus for the stone dropping task, the data shows a combination of pre-test and critical test trial data if subjects did not succeed until the critical test. The combination data (the two graphs on the right) shows the amount of time the subjects were specifically touching the apparatus with a tool being held in their beak.

After subjects’ first successful stone drop, their latency to drop a stone in the following trials greatly decreased (Figure 4, left). Additionally, all subjects began to drop more than one stone in the trials following their first success (Figure 4, right; Table 2; supplementary video 4). Subjects also showed a slight increase in the number of stones they placed on top of the apparatus, but not inside the apparatus, on the trials after the first stone drop (Figure 5).

**Table 2.**
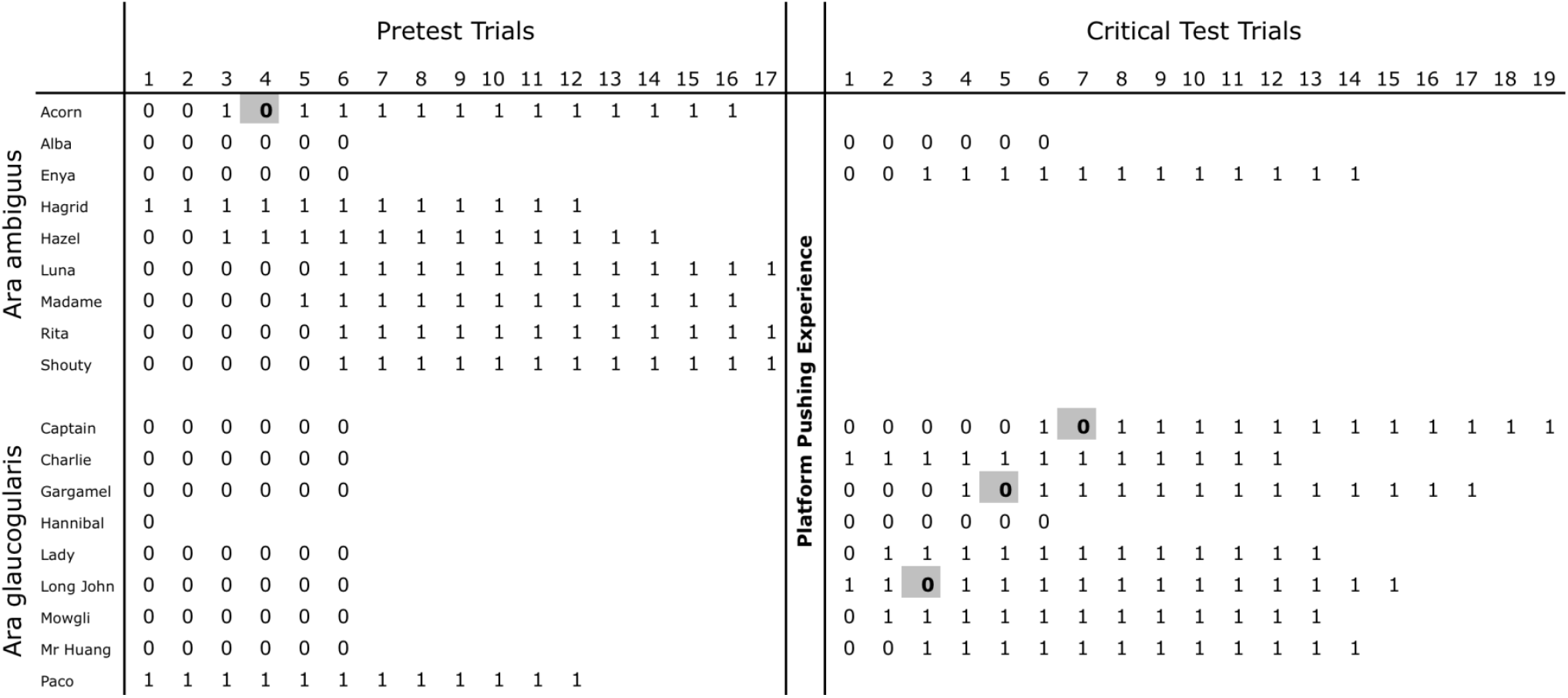
The subjects success across trials in the experiment, showing which subjects succeeded in the pre-test trials and which succeeded in the critical test trials. 0 denotes a failed trial and 1 denotes a successful trial. Trials in which subjects failed after a successful trial are highlighted. Hannibal only recorded one valid pre-test trial, however he had 10 invalid pre-test trials, so he moved onto the experience phase regardless.

**Figure 4.**
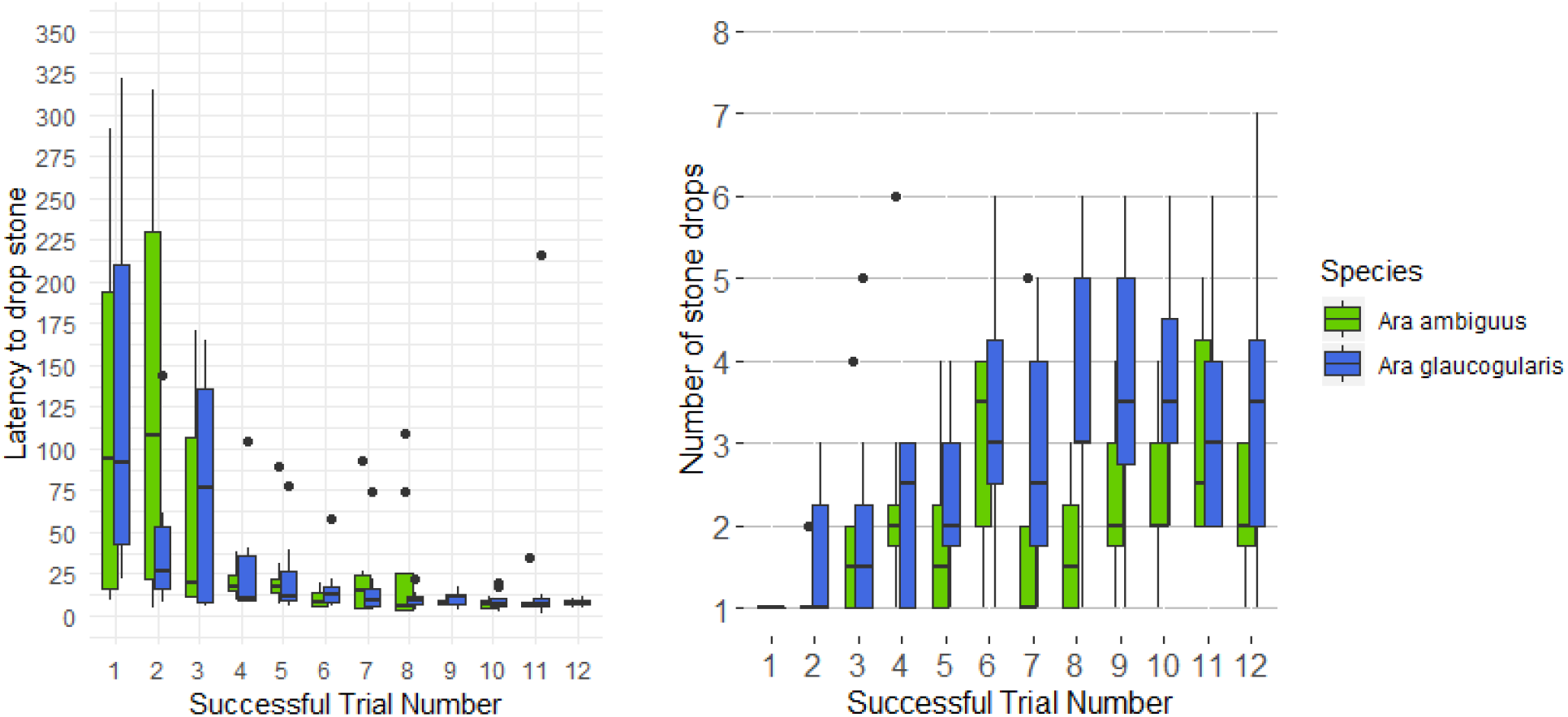
The Subjects behaviour after first success with stone dropping apparatus. After their first successful trial, subjects’ latency to repeat the successful stone dropping behaviour reduced in following trials. They also started increasing the number of stones that they dropped into the apparatus within each trial.

**Figure 5.**
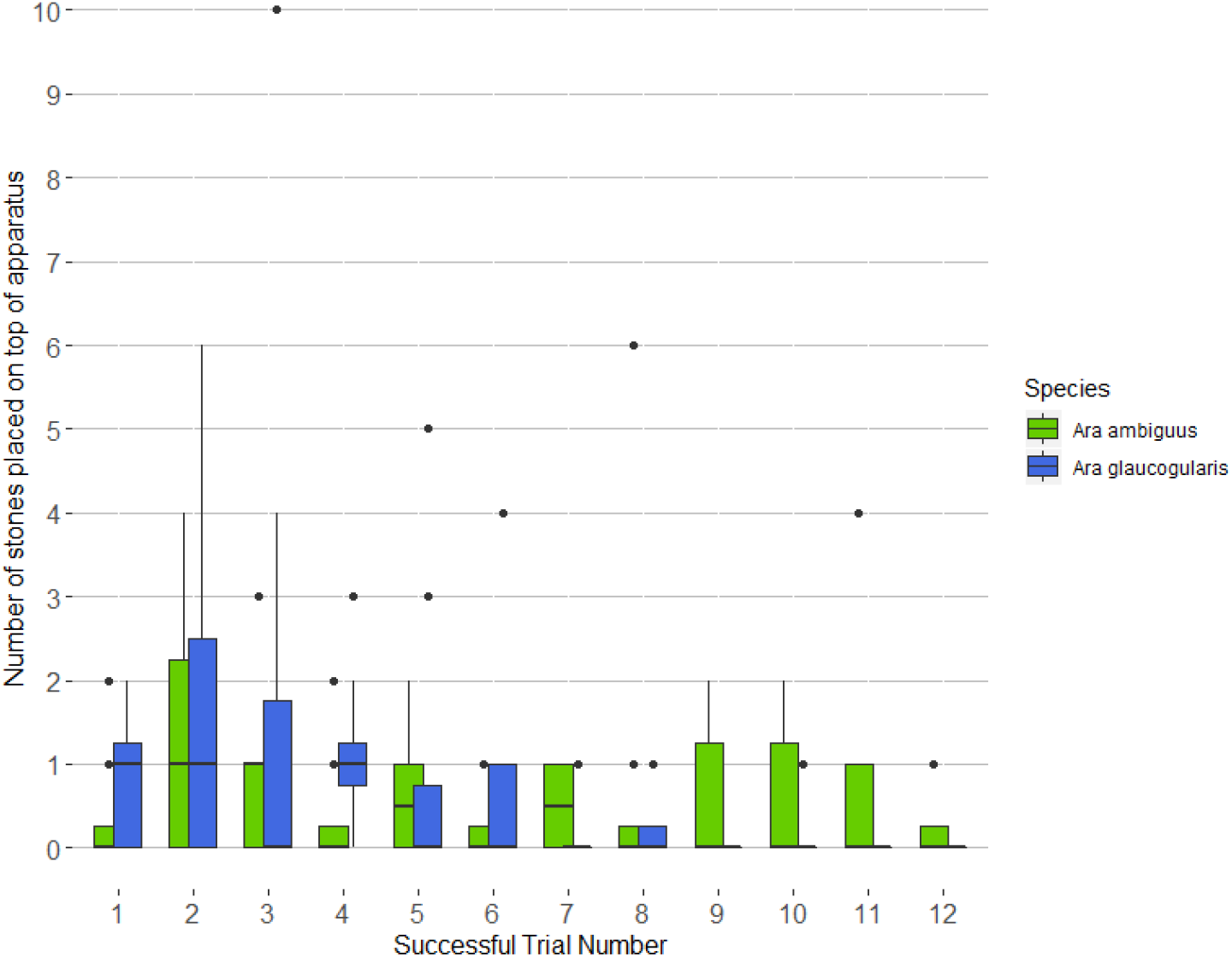
The number of times subjects placed stones on top of the apparatus, but not inside the box. There was a slight peak of placing stones on top of the apparatus, and not inside the box, after the first successful stone-drop. This is suggestive of the subjects recognising they had completed an action that involved the stone and the apparatus, but not specifically placing the stone inside the apparatus.

Only one bird, an *A. glaucogularis* called Mowgli, was able to consistently use the stick tools to make the platform collapse (supplementary video 4). One more *A. glaucogularis*, Charlie, had a single successful trial, but was unable to repeat this in subsequent trials. Four more *A. glaucogularis* inserted the stick tools into the apparatus without hitting the platform and none of the *A. ambiguus* ever inserted a stick tool into the apparatus. Only the two birds that had a successful trial inserted the stick in more than one trial.

In total, the successful bird Mowgli inserted sticks into the apparatus 62 times and obtained the reward from 15 of those cases. Charlie inserted sticks the second most number of times, but he only did so five times, with one successful trial (Table 3).

**Table 3.**
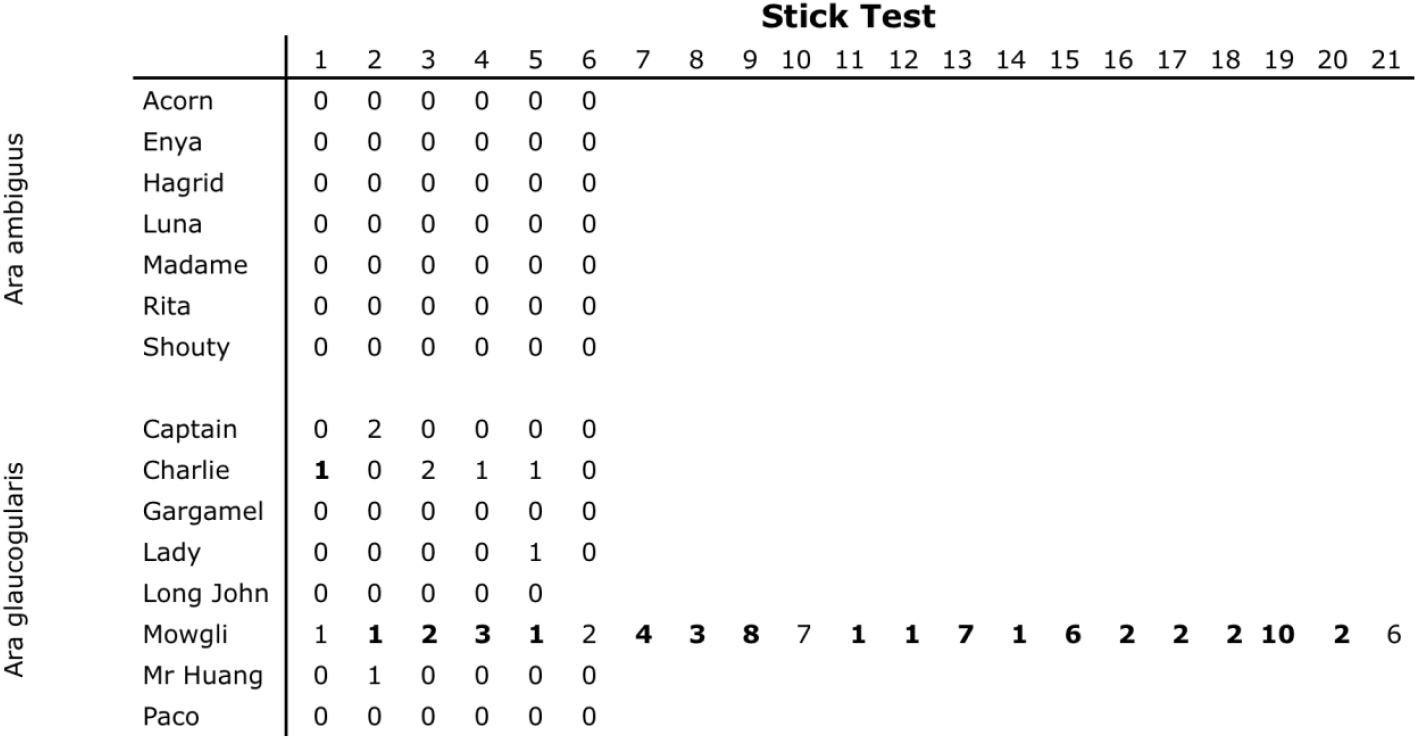
The number of times subjects inserted the stick into the tube in the stick test transfer. Trials in which subjects succeeded to make the platform collapse are in **Bold**. Mowgli did not reach the success criteria of ‘12 uninterrupted successful trials without a failure inbetween’, but we stopped as he was becoming less motivated to participate in the experiment. There was a video error in Long johns sixth trial, hence it is not recorded here.

|  |  | Stick Test |  |  |  |  |  |  |  |  |  |  |  |  |  |  |  |  |  |  |  |  |
| --- | --- | --- | --- | --- | --- | --- | --- | --- | --- | --- | --- | --- | --- | --- | --- | --- | --- | --- | --- | --- | --- | --- |
|  |  | 1 | 2 | 3 | 4 | 5 | 6 | 7 | 8 | 9 | 10 | 11 | 12 | 13 | 14 | 15 | 16 | 17 | 18 | 19 | 20 | 21 |
| Ara ambiguus | Acorn | 0 | 0 | 0 | 0 | 0 | 0 |  |  |  |  |  |  |  |  |  |  |  |  |  |  |  |
|  | Enya | 0 | 0 | 0 | 0 | 0 | 0 |  |  |  |  |  |  |  |  |  |  |  |  |  |  |  |
|  | Hagrid | 0 | 0 | 0 | 0 | 0 | 0 |  |  |  |  |  |  |  |  |  |  |  |  |  |  |  |
|  | Luna | 0 | 0 | 0 | 0 | 0 | 0 |  |  |  |  |  |  |  |  |  |  |  |  |  |  |  |
|  | Madame | 0 | 0 | 0 | 0 | 0 | 0 |  |  |  |  |  |  |  |  |  |  |  |  |  |  |  |
|  | Rita | 0 | 0 | 0 | 0 | 0 | 0 |  |  |  |  |  |  |  |  |  |  |  |  |  |  |  |
|  | Shouty | 0 | 0 | 0 | 0 | 0 | 0 |  |  |  |  |  |  |  |  |  |  |  |  |  |  |  |
| Ara glaucogularis | Captain | 0 | 2 | 0 | 0 | 0 | 0 |  |  |  |  |  |  |  |  |  |  |  |  |  |  |  |
|  | Charlie | **1** | 0 | 2 | 1 | 1 | 0 |  |  |  |  |  |  |  |  |  |  |  |  |  |  |  |
|  | Gargamel | 0 | 0 | 0 | 0 | 0 | 0 |  |  |  |  |  |  |  |  |  |  |  |  |  |  |  |
|  | Lady | 0 | 0 | 0 | 0 | 1 | 0 |  |  |  |  |  |  |  |  |  |  |  |  |  |  |  |
|  | Long John | 0 | 0 | 0 | 0 | 0 |  |  |  |  |  |  |  |  |  |  |  |  |  |  |  |  |
|  | Mowgli | 1 | **1** | **2** | **3** | **1** | 2 | **4** | **3** | **8** | 7 | **1** | **1** | **7** | **1** | **6** | **2** | **2** | **2** | **10** | **2** | 6 |
|  | Mr Huang | 0 | 1 | 0 | 0 | 0 | 0 |  |  |  |  |  |  |  |  |  |  |  |  |  |  |  |
|  | Paco | 0 | 0 | 0 | 0 | 0 | 0 |  |  |  |  |  |  |  |  |  |  |  |  |  |  |  |

There was no difference in exploration rates either between the two species in the task (Figure 3). One *A. ambiguus*, Hazel, did not record a single valid trial as she learned how to break the apparatus

## Discussion

16 out of 18 of parrots of both species were successful in innovating a novel tool use behaviour, i.e. stone-dropping, to solve a novel task (Table 1). The novel stone-dropping behaviour was within the behavioural capacity of both, however, there was a distinct difference between the two species. Seven out of eight of the successful *Ara ambiguus*, but only one *Ara glaucogularis*, solved the task in the pre-test phase, before they received the platform pushing experience, i.e. before they had learned anything about the causal mechanism of the collapsibility of the platform in the apparatus. However, seven out of the eight successful *Ara glaucogularis* and one *Ara ambiguus* solved the task only in the subsequent critical test phase, thus only after they had obtained information about the causal mechanism in the apparatus from the platform pushing experience (Figure 2). The different stages of the experiment at which the solution was innovated suggest that i) there were differences both between and within the species in what types of cognition they used to come up with the behavioural innovation and ii) the stone dropping test alone may not be diagnostic of the types of cognition used for the initial behavioural innovation, which will be discussed further in turn below. As almost all individuals were able to replicate the stone-dropping innovation after their initial success, it suggests that all the subjects at least understood the difference making causal effects of this behaviour. Moreover, the almost uniform failure of the subjects in the stick-test transfer task suggests that their understanding of the platform’s collapsibility was inflexible and did not rely on an understanding of the mechanism. They were mostly unable to redirect a stick tool towards a platform in a different position. Nevertheless, a single individual was able to consistently succeed in this transfer task, showing that there are some conditions that may encourage these macaws to seek the underlying causal mechanism of an apparatus.

The original aim of this experiment was to examine whether an understanding of the functional mechanism of the task could underlie a subjects’ problem solving performance in the stone dropping apparatus (Bird and Emery, 2009; von Bayern et al., 2009). In total, eight parrots (7 *Ara glaucogularis*, and 1 *Ara ambiguus*, Figure 2) were able to innovate the stone dropping behaviour after the platform pushing experience, similar to two out of four New Caledonian crows (von Bayern et al., 2009). The successful innovation of the behaviour at this stage suggests that the subjects had used a mechanistic understanding of how the platform required some form of force or contact to collapse, and had therefore dropped the stone onto it to recreate the force/contact they had previously exerted with their beaks.

The other eight successful parrots (7 *Ara ambiguus* and 1 *Ara glaucogularis*) were all successful in the pre-test phase of the experiment (Figure 2), prior to having any functional experience of the collapsibility of the platform within the apparatus. This was unexpected as it did not happen in the first iteration of this experiment with New Caledonian crows (von Bayern et al., 2009). At this stage, it was impossible for the parrots to be aware of the mechanics of the task because they had never observed the platform moving. For these parrots, the initial innovation of the solution was without a causal understanding of how the dropping stone would affect the platform; they could not have predicted the effect the stone would have. It is likely that these eight individuals innovated their solution through exploratory behaviour.

This would suggest that *Ara glaucogularis* are more likely to need mechanistic information when compared to the the *Ara ambiguus* to innovate the stone-dropping task solution and it may be that the *Ara glaucogularis*’ causal understanding is more biased towards causal mechanisms (Johnson and Ahn, 2017). As the majority of the *Ara ambiguus* solved the task before getting to the mechanism information stage (the platform pushing), it would suggest that they are perhaps more exploratory innovators. Nevertheless, a single individual from both species solved the task in ‘non species typical’ way. This suggests that there are also individual differences within the species, as one *Ara glaucogularis* appears to be an ‘exploratory innovator’ and one *Ara ambiguus* appears to be a ‘mechanistic innovator’. Both species appear to have some flexibility in their innovation types. Furthermore, there was one individual of both species that failed to innovate a solution at any stage, showing ‘non-innovators’ may also exist within both species (Table 2).

As stated, it was unexpected for the subjects to innovate a solution in the pre-test phase of the experiment. For these individuals, it may be that they arrived at the solution through increased intensity of exploring both the apparatus and tools available to them and serendipitously discovered a beneficial object combination, namely placing the stone into the tube and dropping it. Within the two species studied in this experiment the *Ara ambiguus* spent significantly more time interacting with the stone tools than the *Ara glaucogularis* (Figure 3), which may explain why the majority of the *Ara ambiguus* innovated stone-dropping in the pre-test phase. There was no difference between the species in the other exploration measures we took, regarding how much the individuals interacted with the apparatus both with their beak and with the stone tools (Figure 3; Table 1). Also, based on these exploratory measures, the single *Ara glaucogularis* (Paco) that innovated the solution in the pre-test was not *more* exploratory than his species peers, and the single *Ara ambiguus* (Enya) to *not* innovate the solution in the pre-test was not *less* exploratory than her species peers (Table 1). Thus random exploration may not be the explanatory factor for the individual differences in solution time. It could be that the exploration was more directed, i.e. the subjects were given a stone and an apparatus with a tube on the top and there was a fairly limited number of exploratory actions they could have taken and placing the stone in the tube was one of them.

Highly explorative individuals have been shown to do better in problem solving tasks previously (Overington et al., 2011). Macaws also have a diverse range of manipulative skills (e.g. Villegas-Retana and Araya-H., 2017) like other parrots (Huber and Gajdon, 2006), which are also important in creating novel exploration types, another factor vital in problem solving (Griffin and Guez, 2014). Different species of parrots are thought to have different rates of exploration based on their ecological backgrounds (Mettke-Hofmann et al., 2002). Although it has been previously found that one macaw species, the northern red-shouldered macaw (*Diopsittaca nobilis*), shows reduced object-exploration intensity compared to New-Caledonian crows (Auersperg et al., 2015), both species of macaws in this study appeared very exploratory. Kea, another parrot species in contrast have been shown to be much more explorative than New Caledonian crows in regards to haptic exploration (Auersperg et al., 2011), so it might be that the *Ara ambiguus* from this study share some ecological factors with the kea that select for increased haptic exploration and a tendency for creating object combinations, which most likely explains why they produced the stone-dropping behaviour in pre-tests in contrast to the New Caledonian crows (von Bayern et al., 2009). In corvids, one raven has previously been noted to arrive at the solution of the stone dropping task before having any experience of the platform collapsing before (Kabadayi and Osvath, 2017). Although these are typically neophobic birds (Heinrich, 1988) they are still exploratory once they have overcome their neophobia (O’Hara et al., 2017).

In order to obtain a direct comparison of the object exploration tendencies of the two macaw species tested here with that of the previously tested New Caledonian crows, ravens and other birds, it would be informative to study their object-combination tendencies as described in Auersperg *et al*. (2015). We do not know much about their ecology in the wild nor their drive to explore, but it appears that the *Ara glaucogularis* are fairly reliant on a single food source (palm fruits) (Collar et al., 2019b; Yamashita and Machado de Barros, 1997) whereas *Ara ambiguus* are forced to search for variable food sources for two months of the year when their preferred food source (mountain almond trees) is not in season (Berg et al., 2007; Collar et al., 2019a). This may have selected for an increased exploratory tendency in *Ara ambiguus* compared to *Ara glaucogularis* and may explain why more of the *Ara ambiguus* discovered the stone dropping solution during the pre-test stage.

If some subjects, regardless of species, were able to innovate the stone-dropping behaviour without the platform-pushing experience, then it is possible that the subjects that required the platform-pushing experience did not specifically gain mechanistic information from the platform-pushing. Rather than learning something about the mechanism during this phase, the platform pushing may have served to more simply create a positive valence for the whole apparatus, i.e. the subjects recognised it was a potentially reward giving object. This could have encouraged the subjects to explore it further, and thus it may have been exploration that ultimately led them to place stones into the tube, not a mechanism based causal understanding. Thus, the subjects that discovered the solution after the platform-pushing experience may have also discovered the solution by chance, but just taking more time as they were less explorative than the subjects that discovered the solution before the platform pushing. To control for this possibility, it would be good to add another control group that instead of the platform pushing experience were given a similar number of sessions to consume rewards from the apparatus at the same phase of the experiment. If this control group also innovate the stone-dropping, then it suggests that there was no mechanistic understanding involved, but a local enhancement of the apparatus encouraging exploration.

Notably, regardless of the cognitive processes behind the innovation, all individuals across both species appeared to recognise and remember which aspect of their own behaviour had led to the reward from the moment they had first solved the task i.e. dropped a stone in the tube for the very first time. All birds that innovated once were able to consistently repeat the behaviour (Table 2). Four birds had a single failed trial after their first success, and the rest had zero failed trials (Table 2). Additionally, the speed with which the behaviour was repeated suggests rapid learning (Figure 4). This strongly suggests that the parrots recognised which of their own actions had influenced their environment, even if their initial behaviour had come from accidental exploration. They recognised the difference making properties this tool had had on the apparatus (Woodward, 2011). However, after the subjects’ first successful stone-drop, there was also a small peak in erroneous stone placement behaviour in some subjects (Figure 5). The subjects increased the number of times they placed stones on top of the apparatus, but not inside the open hole to the platform, in the trials immediately after their first success. This suggests that the subjects perhaps only recognised a more general behaviour involving both the stone and the apparatus was important to creating the desired effect and still needed some trial and error before recognising the exact behaviour of placing the stone inside the open hole above the platform.

Further evidence that the subjects perhaps did not fully understand the mechanism of the apparatus comes from the almost uniform failure in the stick test transfer. Only five of the subjects inserted the sticks into the apparatus, all of them were *Ara glaucogularis* and only two did so in more than one trial (Table 3). The majority of subjects did not appear to be able to flexibly transfer their knowledge of dropping stones into the apparatus to dropping a stick into the apparatus. Most of the subjects that did insert a stick into the apparatus did not do so persistently as for most of them it did not lead to the platform collapsing, as the platform had now been moved from directly underneath the tube. Interestingly, none of the subjects that solved the stone dropping task in the pre-test ever inserted a stick into the apparatus. It could be that those that required knowledge of the mechanism to solve the task gained a more flexible causal understanding of the apparatus, hence they were more flexible in what kind of tool to try and insert into the apparatus. However, based on personal observations of the stick-test, it was apparent that the subjects struggled to handle the stick tools, and this may have been the underlying factor for why most subjects did not insert them into the tube.

The single subject that did succeed in multiple trials, Mowgli, was not very consistent in producing the successful behaviour. He required many attempted insertions and many failed ‘drops’ of the stick tool before he could make it consistently come into contact with the platform (see supplementary video 4 for an example). Previous research has shown that persistence is a key factor in problem-solving tasks (Chow et al., 2016; Thornton and Samson, 2012), and it is presumably Mowgli’s persistence that allowed him to succeed as he was the only one to repeatedly attempt to use the stick tool (Table 3). None of the macaws were able to use the sticks as a skilled tool user would: by holding one end of it and directing the distal end of the stick in the desired direction. Instead, they used it as an awkwardly shaped stone, dropping the proximal end they were holding vertically into the tube. This did not give the macaws much control over the stick, ergo it was probably very difficult for them to direct the stick towards the moved platform. Nevertheless, Mowgli’s skill with the stick appeared to improve across the course of his trials and he required fewer attempts to drop the stick onto the platform before succeeding in later trials (Table 3). This would suggest that he had paid attention to the detail that the platform had moved as he was directing the sticks towards the new position of the platform and not simply dropping the stick into the tube undirected. It’s possible he had a mental representation that he had to create contact between the tool and the platform, which in turn suggests he was using a mental representation of part of the causal mechanism of the task. However, Mowgli was the only individual that showed this possibility. Thus the overwhelming evidence from the other subjects is that they do not have a drive to use causal mechanisms to solve problem-solving tasks.

There is other evidence that suggests the subjects appear to have not quite understood exactly how the stone-tool influenced the moveable platform. All of the subjects that started stone dropping also began to drop more than one stone in the following trials (Figure 4, right; see supplementary video 3 for example). This behaviour suggests the subjects did not make the connection that the stone they had dropped had impacted the platform, which in turn made it collapse. Instead, it suggests they may have been trying to get more rewards from an apparatus that had already released its reward. This mistake implies the subjects only recognised that dropping the stone in the hole led to a reward, not how or why it did. This behaviour could also be explained by a lack of inhibitory control as it had become a conditioned behaviour (Kabadayi et al., 2017). Equally, the repeated stone dropping may have been additional play and exploratory behaviour, which as discussed above, could be the reason they found the solution in the first place, and probably a hard behaviour to suppress.

In conclusion, there is strong evidence that these macaws are good problem-solvers as the majority were able to innovate a tool-using solution to the stone-dropping task, i.e. they were able to drop a stone onto a collapsible platform. There is some evidence to suggest that some subjects, mostly *Ara glaucogularis*, used a causal understanding of the mechanism underlying the apparatus, the collapsible platform, to innovate the initial stone-dropping solution. There is also evidence to suggest that most of the *Ara ambiguus* used exploratory behaviour to discover the stone-dropping innovation. Nevertheless, all subjects displayed an understanding of which of their behaviours had been critical in making the platform collapse as all of them were able to rapidly and consistently repeat their stone-dropping innovations. Thus they all displayed a difference-making, learned, causal understanding of their behaviour. Most of the subjects had difficulties in handling stick tools in the transfer task, but one *Ara glaucogularis* individual was able to succeed in this task, which gave stronger evidence that he had used an understanding of the underlying mechanism to innovate the use of the tools to collapse the platform.

## Supporting information

Raw data file

## Ethical Statement

All applicable international, national, and institutional guidelines for the care and use of animals were followed. In accordance with the German Animal Welfare Act of 25th May 1998, Section V, Article 7 and the Spanish Animal Welfare Act 32/2007 of 7th November 2007, Preliminary Title, Article 3, the study was classified as non-animal experiment and did not require any approval from a relevant body.

## Supplementary data and videos

Data from the video coding and Supplementary videos can be found on figshare at: https://doi.org/10.6084/m9.figshare.12800768.v1

## Author Contributions

The design of the experiment was initially conceived by LO’N and AvB, with contributions from AP and NB. The experiments were carried out by AP, NB and RH. Videos were coded by RH and data were visualised by LO’N. The initial manuscript was written by LO’N with feedback from AvB and MG.

## Acknowledgements

We thank Magdalena van Buuren & Luisana Carballo for their comments on earlier versions of the manuscript.

We thank the Loro Parque and its president, Mr Wolfgang Kiessling for their support, the access to the birds and the research facilities. We thank the Loro Parque Fundación and its president Mr Christoph Kiessling for their collaboration and the staff of the Loro Parque Fundación, the animal caretakers and the veterinary department for their support.

LO’N is a member of the International Max Planck Research School (IMPRS) for Organismal Biology. The study was funded by the Max-Planck Society.

## Notes

https://doi.org/10.6084/m9.figshare.12800768.v1

### Competing Interest Statement

The authors have declared no competing interest.

## References

Auersperg, A., von Bayern, A., Gajdon, G., Huber, L., Kacelnik, A., 2011. Flexibility in Problem Solving and Tool Use of Kea and New Caledonian Crows in a Multi Access Box Paradigm. PLoS ONE 6, e20231. https://doi.org/10.1371/journal.pone.0020231

Auersperg, A.M.I., Huber, L., Gajdon, G.K., 2011. Navigating a tool end in a specific direction: stick-tool use in kea (Nestor notabilis). Biol. Lett. 7, 825–828. https://doi.org/10.1098/rsbl.2011.0388

Auersperg, A.M.I., Szabo, B., von Bayern, A.M.P., Kacelnik, A., 2012. Spontaneous innovation in tool manufacture and use in a Goffin’s cockatoo. Curr. Biol. 22, R903–R904. https://doi.org/10.1016/j.cub.2012.09.002

Auersperg, A.M.I., van Horik, J.O., Bugnyar, T., Kacelnik, A., Emery, N.J., von Bayern, A.M.P., 2015. Combinatory actions during object play in psittaciformes (Diopsittaca nobilis, Pionites melanocephala, Cacatua goffini) and corvids (Corvus corax, C. monedula, C. moneduloides). J. Comp. Psychol. 129, 62–71. https://doi.org/10.1037/a0038314

Berg, K.S., Socola, J., Angel, R.R., 2007. Great Green Macaws and the annual cycle of their food plants in Ecuador. J. Field Ornithol. 78, 1–10. https://doi.org/10.1111/j.1557-9263.2006.00080.x

Bird, C.D., Emery, N.J., 2009. Insightful problem solving and creative tool modification by captive nontool-using rooks. Proc. Natl. Acad. Sci. 106, 10370–10375. https://doi.org/10.1073/pnas.0901008106

Chow, P.K.Y., Lea, S.E.G., Leaver, L.A., 2016. How practice makes perfect: the role of persistence, flexibility and learning in problem-solving efficiency. Anim. Behav. 112, 273–283. https://doi.org/10.1016/j.anbehav.2015.11.014

Collar, N., Boesman, P., Sharpe, C.J., 2019a. Great Green Macaw (Ara ambiguus).

Collar, N., Boesman, P., Sharpe, C.J., del Hoyo, J., Elliott, A., Sargatal, J., Christie, D.A., de Juana, E., 2019b. Blue-throated Macaw (Ara glaucogularis), in: Handbook of the Birds of the World Alive. Lynx Editions.

Emery, N.J., Clayton, N.S., 2004. The Mentality of Crows: Convergent Evolution of Intelligence in Corvids and Apes. Science 306, 1903–1907. https://doi.org/10.1126/science.1098410

Goodman, M., Hayward, T., Hunt, G.R., 2018. Habitual tool use innovated by free-living New Zealand kea. Sci. Rep. 8, 13935. https://doi.org/10.1038/s41598-018-32363-9

Griffin, A.S., Guez, D., 2014. Innovation and problem solving: A review of common mechanisms. Behav. Processes 109, 121–134. https://doi.org/10.1016/j.beproc.2014.08.027

Gutiérrez-Ibáñez, C., Iwaniuk, A.N., Wylie, D.R., 2018. Parrots have evolved a primate-like telencephalic-midbrain-cerebellar circuit. Sci. Rep. 8, 9960. https://doi.org/10.1038/s41598-018-28301-4

Hanus, D., 2016. Causal reasoning versus associative learning: A useful dichotomy or a strawman battle in comparative psychology? J. Comp. Psychol. 130, 241–248. https://doi.org/10.1037/a0040235

Heinrich, B., 1988. Why Do Ravens Fear Their Food? The Condor 90, 950–952. https://doi.org/10.2307/1368859

Heinsohn, R., Zdenek, C.N., Cunningham, R.B., Endler, J.A., Langmore, N.E., 2017. Tool-assisted rhythmic drumming in palm cockatoos shares key elements of human instrumental music. Sci. Adv. 3, e1602399. https://doi.org/10.1126/sciadv.1602399

Herculano-Houzel, S., 2017. Numbers of neurons as biological correlates of cognitive capability. Curr. Opin. Behav. Sci. 16, 1–7. https://doi.org/10.1016/j.cobeha.2017.02.004

Huber, L., Gajdon, G.K., 2006. Technical intelligence in animals: the kea model. Anim. Cogn. 9, 295–305. https://doi.org/10.1007/s10071-006-0033-8

Iwaniuk, A.N., Dean, K.M., Nelson, J.E., 2005. Interspecific Allometry of the Brain and Brain Regions in Parrots (Psittaciformes): Comparisons with Other Birds and Primates. Brain. Behav. Evol. 65, 40–59. https://doi.org/10.1159/000081110

Johnson, S.G.B., Ahn, W., 2017. Causal Mechanisms. Oxford University Press. https://doi.org/10.1093/oxfordhb/9780199399550.013.12

Kabadayi, C., Krasheninnikova, A., O’Neill, L., van de Weijer, J., Osvath, M., von Bayern, A.M.P., 2017. Are parrots poor at motor self-regulation or is the cylinder task poor at measuring it? Anim. Cogn. 20, 1137–1146. https://doi.org/10.1007/s10071-017-1131-5

Kabadayi, C., Osvath, M., 2017. Ravens parallel great apes in flexible planning for tool-use and bartering. Science 357, 202–204. https://doi.org/10.1126/science.aam8138

Kacelnik, A., 2009. Tools for thought or thoughts for tools? Proc. Natl. Acad. Sci. 106, 10071–10072. https://doi.org/10.1073/pnas.0904735106

Krasheninnikova, A., Berardi, R., Lind, M.-A., O’Neill, L., von Bayern, A.M.P., 2019. Primate cognition test battery in parrots. Behaviour 1–41. https://doi.org/10.1163/1568539X-0003549

Kumar, S., Stecher, G., Suleski, M., Hedges, S.B., 2017. TimeTree: A Resource for Timelines, Timetrees, and Divergence Times. Mol. Biol. Evol. 34, 1812–1819. https://doi.org/10.1093/molbev/msx116

Lambert, M.L., Jacobs, I., Osvath, M., von Bayern, A.M.P., 2018. Birds of a feather? Parrot and corvid cognition compared. Behaviour 1–90. https://doi.org/10.1163/1568539X-00003527

Le Pelley, M.E., Griffiths, O., Beesley, T., 2017. Associative Accounts of Causal Cognition. Oxford University Press. https://doi.org/10.1093/oxfordhb/9780199399550.013.2

Lefebvre, L., Whittle, P., Lascaris, E., Finkelstein, A., 1997. Feeding innovations and forebrain size in birds. Anim. Behav. 53, 549–560. https://doi.org/10.1006/anbe.1996.0330

Mettke-Hofmann, C., Winkler, H., Leisler, B., 2002. The Significance of Ecological Factors for Exploration and Neophobia in Parrots. Ethology 108, 249–272. https://doi.org/10.1046/j.1439-0310.2002.00773.x

Neilands, P.D., Jelbert, S.A., Breen, A.J., Schiestl, M., Taylor, A.H., 2016. How Insightful Is ‘Insight’? New Caledonian Crows Do Not Attend to Object Weight during Spontaneous Stone Dropping. PLOS ONE 11, e0167419. https://doi.org/10.1371/journal.pone.0167419

O’Hara, M., Mioduszewska, B., von Bayern, A., Auersperg, A., Bugnyar, T., Wilkinson, A., Huber, L., Gajdon, G.K., 2017. The temporal dependence of exploration on neotic style in birds. Sci. Rep. 7, 4742. https://doi.org/10.1038/s41598-017-04751-0

Olkowicz, S., Kocourek, M., Lučan, R.K., Porteš, M., Fitch, W.T., Herculano-Houzel, S., Němec, P., 2016. Birds have primate-like numbers of neurons in the forebrain. Proc. Natl. Acad. Sci. 113, 7255–7260. https://doi.org/10.1073/pnas.1517131113

O’Neill, L., Picaud, A., Maehner, J., Gahr, M., von Bayern, A.M.P., 2018. Two macaw species can learn to solve an optimised two-trap problem, but without functional causal understanding. Behaviour 1–30. https://doi.org/10.1163/1568539X-00003521

Osuna-Mascaró, A.J., Auersperg, A.M.I., 2018. On the brink of tool use? Could object combinations during foraging in a feral Goffin’s cockatoo (Cacatua goffiniana) result in tool innovations? Anim. Behav. Cogn. 5, 229–234. https://doi.org/10.26451/abc.05.02.05.2018

Osvath, M., Kabadayi, C., Jacobs, I., 2014. Independent Evolution of Similar Complex Cognitive Skills: The Importance of Embodied Degrees of Freedom. Anim. Behav. Cogn. 1, 249. https://doi.org/10.12966/abc.08.03.2014

Overington, S.E., Cauchard, L., Côté, K.-A., Lefebvre, L., 2011. Innovative foraging behaviour in birds: What characterizes an innovator? Behav. Processes 87, 274–285. https://doi.org/10.1016/j.beproc.2011.06.002

Povinelli, D.J., 2000. Folk physics for apes: the chimpanzee’s theory of how the world works, Dig. print. (repr. 2008). ed. Oxford Univ. Press, Oxford.

Sol, D., Duncan, R.P., Blackburn, T.M., Cassey, P., Lefebvre, L., 2005. Big brains, enhanced cognition, and response of birds to novel environments. Proc. Natl. Acad. Sci. 102, 5460–5465. https://doi.org/10.1073/pnas.0408145102

Sol, D., Székely, T., Liker, A., Lefebvre, L., 2007. Big-brained birds survive better in nature. Proc. R. Soc. B Biol. Sci. 274, 763–769. https://doi.org/10.1098/rspb.2006.3765

Sol, D., Timmermans, S., Lefebvre, L., 2002. Behavioural flexibility and invasion success in birds. Anim. Behav. 63, 495–502. https://doi.org/10.1006/anbe.2001.1953

Thornton, A., Samson, J., 2012. Innovative problem solving in wild meerkats. Anim. Behav. 83, 1459–1468. https://doi.org/10.1016/j.anbehav.2012.03.018

Toft, C.A., Wright, T.F., Gilardi, J.D., 2016. Parrots of the wild: a natural history of the world’s most captivating birds.

Van Horik, J.O., Clayton, N.S., Emery, N.J., 2012. Convergent Evolution of Cognition in Corvids, Apes and Other Animals. Oxford University Press. https://doi.org/10.1093/oxfordhb/9780199738182.013.0005

Villegas-Retana, S.A., Araya-H., D., 2017. Consumo de almendro de playa (Terminalia catappa) y uso de hojas como herramienta por parte del ave Ara ambiguus (Psittaciformes: Psittacidae) en Costa Rica. UNED Res. J. 9. https://doi.org/10.22458/urj.v9i2.1894

von Bayern, A.M.P., Heathcote, R.J.P., Rutz, C., Kacelnik, A., 2009. The Role of Experience in Problem Solving and Innovative Tool Use in Crows. Curr. Biol. 19, 1965–1968. https://doi.org/10.1016/j.cub.2009.10.037

Wood, G.A., 1984. Tool use by the Palm Cockatoo Probosciger aterrimus during display. Corella 8, 94–95.

Woodward, J., 2011. A Philosopher Looks at Tool Use and Causal Understanding, in: Tool Use and Causal Cognition. Oxford University Press.

Wright, T.F., Schirtzinger, E.E., Matsumoto, T., Eberhard, J.R., Graves, G.R., Sanchez, J.J., Capelli, S., Müller, H., Scharpegge, J., Chambers, G.K., Fleischer, R.C., 2008. A Multilocus Molecular Phylogeny of the Parrots (Psittaciformes): Support for a Gondwanan Origin during the Cretaceous. Mol. Biol. Evol. 25, 2141–2156. https://doi.org/10.1093/molbev/msn160

Yamashita, C., Machado de Barros, Y., 1997. The Blue-throated Macaw Ara glaucogularis: characterisation of its distinctive habitats in savannahs of the Beni, Bolivia. Ararajuba 5, 141–150.

